## Supplemental Figures for "Population Genomics of *Plasmodium vivax* in Panama to Assess the Risk of Case Importation on Malaria Elimination"

**Supporting information**

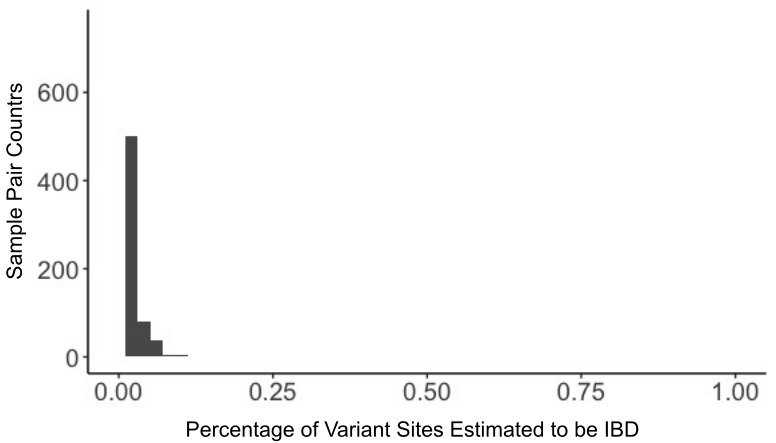

**S1 Figure. Distribution of IBD in Panamanian-Colombian sample pairs.**

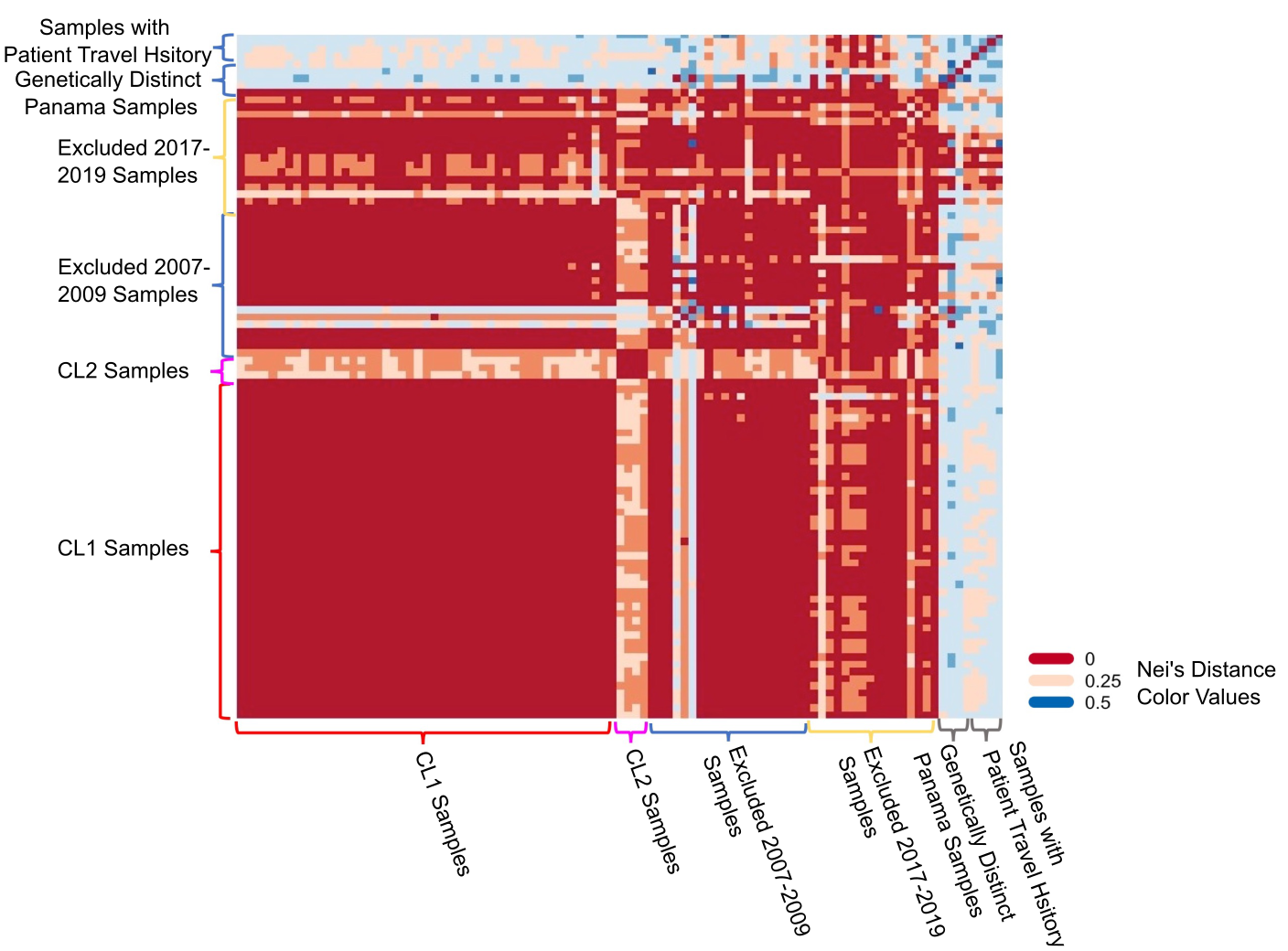

**S2 Figure.** **Annotated heatmap of pairwise Nei’s standard distance comparisons between all 2007-2009 and 2017-2019 samples using SNPs that were callable in at least 80% of samples. Each block row and column presents a single sample**. Brackets indicate sample groups.

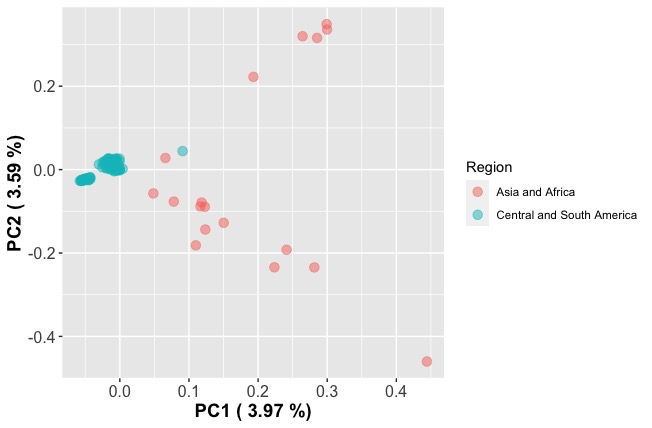

**S3 Figure.** **Principal components analysis of Panama samples and previously collected samples from Central and South America, Asia, and Africa.** Samples are colored by the region of origin. Parentheses contain the percentage of variance explained by each principal component.

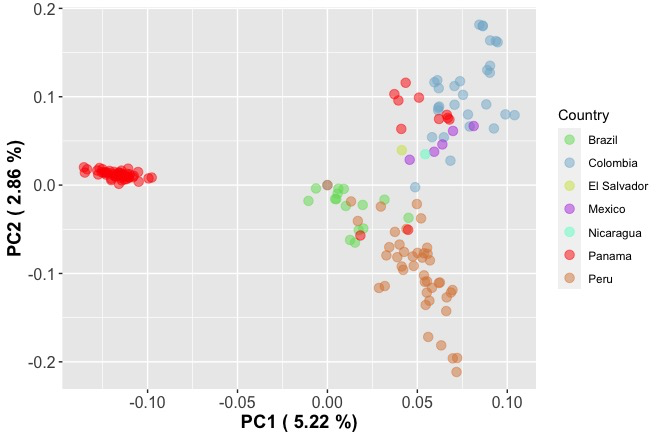

**S4 Figure.** **Principal components analysis of Panama samples and previously collected Central and South American samples.** Samples are colored by country of origin. Parentheses contain the percentage of variance explained by each principal component.
